# A novel data-driven model for real-time influenza forecasting

**DOI:** 10.1101/185512

**Authors:** Siva R. Venna, Amirhossein Tavanaei, Raju N. Gottumukkala, Vijay V. Raghavan, Anthony Maida, Stephen Nichols

## Abstract

We provide data-driven machine learning methods that are capable of making real-time influenza forecasts that integrate the impacts of climatic factors and geographical proximity to achieve better forecasting performance. The key contributions of our approach are both applying deep learning methods and incorporation of environmental and spatio-temporal factors to improve the performance of the influenza forecasting models. We evaluate the method on Influenza Like Illness (ILI) counts and climatic data, both publicly available data sets. Our proposed method outperforms existing known influenza forecasting methods in terms of their Mean Absolute Percentage Error and Root Mean Square Error. The key advantages of the proposed data-driven methods are as following: (1) The deep-learning model was able to effectively capture the temporal dynamics of flu spread in different geographical regions, (2) The extensions to the deep-learning model capture the influence of external variables that include the geographical proximity and climatic variables such as humidity, temperature, precipitation and sun exposure in future stages, (3) The model consistently performs well for both the city scale and the regional scale on the Google Flu Trends (GFT) and Center for Disease Control (CDC) flu counts. The results offer a promising direction in terms of both data-driven forecasting methods and capturing the influence of spatio-temporal and environmental factors for influenza forecasting methods.

## I. Introduction

Seasonal influenza is a major global health issue that affects many people across the world. According to the Center for Disease Control (CDC) reports [1] in the United States alone there were 9.2 million to 60.8 million reported illnesses since 2010. Influenza can cause severe illness and even death for high risk populations. For instance, during 2012-2013-which was a pretty bad flu season-the outbreak has resulted in 56,000 deaths and 710,000 hospitalizations. Prevention and control of influenza spread can be a huge challenge, especially without adequate tools that can monitor and also predict future outbreaks in various populations. With accurate and reliable prediction of influenza outbreaks, public health officials would be able to mitigate the effects of widespread outbreak through aggressive measures, prioritizing resources in terms of staff, vaccines and emergency rooms to prevent widespread outbreaks. Predicting influenza is a very difficult task given the complicated stochastic characteristics of the influenza strain and environmental conditions that affect the severity of the spread. Given the importance of this problem, many researchers have investigated various aspects of influenza including the dynamics of spread and future forecasting. CDC [2], [3], [4] and Defense Advanced Research Projects Agency (DARPA) [5], [6] have launched several competitions to solve the problem of real-time forecasting of influenza and other infectious diseases. Forecasting influenza remains an active research area given the limited ability of existing models to effectively capture the dynamics of the influenza spread across different populations and environmental conditions while improving the limited accuracy of existing forecasting models.

Influenza forecasting research is broadly classified into three categories. The first category includes traditional compartment models such as Susceptible-Infected-Recovered (SIR) [7], [8], Susceptible-Infected-Recovered-Susceptible (SIRS) [9], [10], and Susceptible-Exposed-Infected-Recovered (SEIR) [11], [12]. The compartmental models are intuitive in terms of capturing the different states of infected populations. These models are deterministic and lack the flexibility to be re-calibrated in terms of capturing the dynamics of influenza spread. The models in the second category employs statistical and time-series based methodologies such as Box-Jenkins, employing some variant of Auto-Regression Integrated Moving Average (ARIMA) [13] and Generalized Autoregressive Moving Average (GARMA) [14]. The Box-Jenkins based time-series methods are flexible in terms of capturing the trending behavior of affected populations, but suffer from poor accuracy as the influence of external factors is not well captured in existing forecasting models. The third category models are machine learning methods that have gained increased prominence in recent years. Some popular machine learning methods include Stacked Linear Regression [15], Support Vector Regression [16], Binomial Chain [17], and Classification and Regression Trees [18]. Machine learning based approaches are data-driven approaches that offer more flexibility in terms of capturing the influence of multiple external variables, but are computationally expensive compared to statistical models, as the model has to be retrained when new data arrives. With advances in computational power, machine learning based models offer a promising direction. Use of machine learning methods in understanding influenza dynamics are discussed in [19], [20], [21]. Additionally, review of existing influenza forecasting methods is discussed in [22], [23], [24].

Recurrent Neural Networks (RNNs) have shown remarkable performance in sequential (temporal) data prediction [25]. However, the conventional RNNs have shown practical difficulties in training the networks faced with long interval temporal contingencies of input/output sequences [26]. Therefore, an efficient gradient-based method called Long Short Term Memory (LSTM) was introduced to develop a stable recurrent architecture [27]. This new technology supersedes RNNs for time series forecasting. In regard to recurrent networks, it solves the vanishing/exploding gradient problem and gives much more flexibility to the learning algorithm on when to forget the past or ignore the current input. This network has been successfully applied to various temporal data processing problems, such as context free/sensitive machine language learning [28], speech processing and recognition [29], [30], and handwriting recognition [31]. An interesting property of this model is the ability of LSTM to learn to selectively forget/remember historical information. The forgetting ability stops the network from growing indefinitely and breaking down [32]. In time series prediction, the few recent value points convey the most relevant information for predicting the future points. The LSTM neural network can also be trained effectively to predict the future points in time series using the few available points [33]. The deep network architecture of the LSTM cells can provide a powerful model in temporal data processing. Recently, LSTM and deep LSTM have attracted much interest in temporal data prediction such as traffic speed prediction [34] and classification of diagnoses given intensive care unit time series [35]. In this paper we explore a deep LSTM neural network for the flu prediction problem. The deep architecture can be fulfilled by unrolling the LSTM cells in which the input of the successor cell is provided by the output of the predecessor cell.

Improving the accuracy of influenza forecasting requires effective integration of external variables that are shown to have strong influence on flu spread. Many traditional and non-traditional data sources have been explored to improve flu forecasting, including: historical Influenza Like Illness (ILI) counts; climate and weather information [14]; social media interactions such as Twitter messages [15], [16] and Google searches involving flu related words [10], Google Flu Trends [14]; and travel patterns [36]. Several environmental factors are known to affect or influence flu counts. These include population size, climate and weather information, travel patterns, infection status in neighborhood cities or regions, rural-urban location differences, etc. Usually ILI or other infectious disease transmission may occur [37], [38], [39], [40] through (1) direct contact with infected subjects, (2) intermediate objects, or (3) droplets and other particles expelled from infected individuals. Previous studies have clearly identified direct influence of weather variables such as temperature, humidity, precipitation etc. on influenza virus transmission and survival [41], [42], [43]. As presented in [42], low relative humidity aids in faster evaporation of expelled droplets or particles and longer survival of the airborne virus. Also, geographical regions that are in close proximity to infected regions have high risk of becoming infected due to population movements and high-likelihood of social interactions [44], [45], [46]. The impact of environmental factors must be integrated effectively into the flu forecasting model to achieve better accuracy with influenza prediction models. Recent work from [14] tried to capture the influence of environmental conditions for flu forecasting using GARMA(3,0) model. Experimental studies in [47], [48], however, demonstrated that temperature and humidity are not linearly correlated with influenza spread. Our work makes a few improvements in terms of how the influence of external environmental variables are captured to further improve the prediction accuracy of our proposed baseline LTSM model. First, we capture situational time lags between the flu counts and the weather variables that produce non-linear correlation. Second, we also capture the influence of the spatial proximity of different geographical regions. We evaluated the model for different spatio-temporal granularity and data sources.

The proposed multi-stage forecasting approach employs an LSTM neural network as a time series forecasting model to forecast influenza counts. The primary contributions of the paper are the introduction of a deep-learning approach to forecast influenza, and a multistage approach to capture the influence of geographical proximity and the impacts of environmental factors. Our proposed method is evaluated on both GFT and CDC data. The LSTM model performs better than the existing baseline time series based ARIMA model. The LSTM model is further improved in terms of its ability to forecast influenza counts at different spatial and temporal scales by capturing both the influence of geographical proximity, and the impacts from environmental factors in future stages.

## II. Materials and methods

The proposed model consists of two stages. In the first stage, a deep learning model based on the LSTM neural network approach is used to estimate initial forecast. In the second stage the error from the initial forecast is reduced by incorporating two different factors: (1) An impact factor obtained from the weather variables (humidity, precipitation, temperature, sun exposure) by extracting situational time lags using symbolic time series approach; and (2) a spatio-temporal adjustment factor obtained by capturing the influence of flu spread from neighbouring regions that are in geographical proximity.

### A. Data description

For influenza activity, two different real-world data sets are chosen. The CDC-reported ILI data for all ten Health and Human Services (HHS) regions between 1997-2016 [1] is the only national level dataset available for the United States. Google Flu Trends (GFT) [49] data (available from 2009 to 2014) is a weekly estimate of influenza activity derived from aggregated search query data. A subset of the GFT dataset including the flu count trends reported for 6 cities from Texas and Louisiana (Austin, Dallas, Houston, San Antonio, Baton Rouge and New Orleans) is selected. The weather data is downloaded from Climate Data Online (CDO) [50], which provides free access to the National Climatic Data Center (NCDC) archive of historical weather and climate data. The weather variables used include precipitation, maximum temperature, minimum temperature, and sun exposure. For each city from the GFT dataset, all available stations from the CDO within that city’s geographical limits are downloaded. For the CDC dataset, all the stations within each HHS region boundary are downloaded from the CDO. The data collected from the CDO for both datasets are then aggregated, for each city or region, by averaging into single weekly summarized time-series. This aggregated data is then cleaned to treat any further missing values, using simple moving average based smoothing. At this time, all of the collected datasets-ILI, GFT and respective weather variables-are weekly summarized time series. For each experiment a combination of training and validation set approach is used, where training and validation sets are in sequence and mutually exclusive. During each training exercise approximately 600 samples are used for training and immediately 20 samples are used for validation with respect to the CDC dataset. At the same time the GFT dataset training and validation sample sizes are approximately 450 and 20 respectively.

### B. Model

The proposed multi-stage forecasting approach includes the following steps. In the first stage, the LSTM neural network is trained on the flu time series of nodes to forecast the initial flu counts. A node refers to a geographical region, which could be a HHS region or a city. In the second stage, the impact of climatic variables and spatio-temporal adjustment factor are added to the flu counts estimated by the LSTM model to reduce the error. The impact component from climatic variables is computed using the time-delayed association analysis between each symbolic time series of weather and flu counts. The spatio-temporal adjustment factor is calculated by averaging the flu variations at nearby data nodes. The proposed models, namely the baseline LSTM model, the LSTM with climatic variable impact (LSTM+CI), and the LSTM with climatic variable impact and spatial adjustment factor (LSTM+CI+SA) are compared with the state-of-the-art ARIMA(3,0,3) model.

1. Deep Long Short Term Memory network:
  a. *LSTM Cell*∷ RNN computes an output sequence (y_1_, y_2_, …, y_*T*_) based on its input sequence (x_1_, x_2_, …, x_*T*_) and its previous state (h_1_, h_2_, …, h_*T*_) as shown in Eq. 1 and Fig. 1.

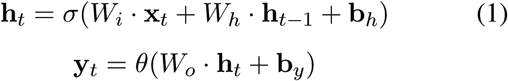

*σ* and *θ* are the hidden and output activation functions. *W* and *b* determine the adaptive weight and bias vectors of the RNN. LSTM is a variation of RNNs preserving back-propagated error through time and layers. Furthermore, the LSTM learning algorithm is local in both space and time, with computational complexity of O(1) per time step and weight [27], which is faster than the popular RNN learning algorithms (e.g. real-time recurrent learning (RTRL) [51] and back-propagation through time (BPTT) [52]). An LSTM cell performs as a memory to write, read, and erase information according to the decisions specified by the input, output, and forget gates, respectively. The weights associated with the gates are trained (adapted) by a recurrent learning process. Fig. 2 shows an LSTM cell containing the input gate, *I*, the forget gate, *F*, and the output gate, *Y*. The memory cell shown in Fig. 2 is implemented as follows:

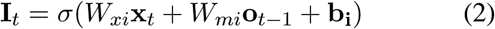

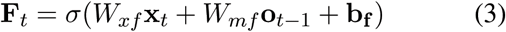

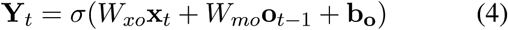

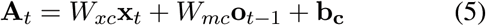

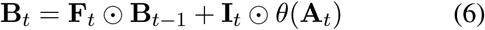

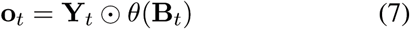

where, *W*_*x*_ and *W*_*m*_ are the adaptive weights, initialized randomly in the range (0,1). x_*t*_ and o_*t*–1_ denote the current input and previous output vectors, respectively. b parameters are bias vectors that are not shown in Fig. 2. The cell state, B_*t*_, is updated by the forget gate, the input gate, and the current input auto-regression value (A_*t*_). σ and *θ* determine the *Sigmoid* and *Tanh* activation functions.
  b. *Deep LSTM Architecture∷* A number of approaches for developing the deep architectures of RNNs and LSTMs have been discussed in the literature [29], [30], [53], [54]. In this investigation, we construct an LSTM network by unrolling the LSTM cells in time. This model provides a suitable architecture for the time series prediction problems due to its sequential framework. Fig. 3 shows the network architecture consisting of the unrolled LSTM cells that are trained by the back propagation algorithm based on the mean-square-error cost function (training criterion). The corresponding LSTM cell at time *t* — *i* receives the flu count calculated by the predecessor cell (*o*_*t*-*i*-1_) and the input, *x*_*t*–*i*_, to calculate the flu count at *t* — *i*, *o*_*t*–*i*_. This process is repeated for all the LSTM cells in the model. The number of LSTM cells denotes the number of time steps, *T*, before the current time. To calculate the flu count at the current state, *t*, the data points from *T* previous time steps are used. After different experimental setups, we selected *T* = 20 time steps.
2. *Climatic Variable Impact*: Each of the climatic variables such as humidity, sun exposure, precipitation, and temperature have different degrees of impact on influenza spread in a geographical region. The impact of these variables has been well studied in the literature [41], [42], [43]. One can observe strong correlation between minimum and maximum temperatures and influenza counts from CDC in Fig. 4. Linear integration of multiple time series is not an effective way to capture the impact of climatic variables because the magnitude and the impact delay of temporal values can change with respect to the geographical location. The composite impact of climatic variables that is added to the original LSTM model is computed by a weighted summation of individual impacts. The overall procedure to obtain the aggregated impact includes (1) Establishing non-linear situational correlation between each weather variable and the flu counts using symbolic time series to obtain situational time lags, and (2) aggregating the individual impacts. To compute the situational time lags between each weather variable and the flu count at a data node, the numerical time-series are converted to symbolic time-series. The symbols at each time step for any variable are shown by a tuple created from the variable set (high, normal, low) and change trend (increasing, stable, decreasing). Once the symbolic time series are generated, time delayed apriori associations are computed in time delays ranging from 0 to 5 weeks, between the flu counts and each weather variable. From these associations, the most confident symbolic pairs for each possible symbol combination are finally selected. Once the time lags between flu counts and each weather variable are computed for all the data nodes, total impact, *I*^tot^, inflicted at time step *t* from the weather variables for data node *n* is estimated using the following formula.

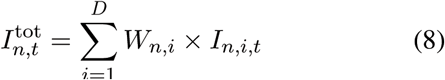 The aggregated impact 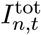 is basically the weighted summation of impact (change) inflicted by D weather variables. The weights, W_*i*_s, are trained using Widrow-Hoff learning [55] with mean square error (MSE) criterion as the cost function on the available data. The target of this Widrow-Hoff learning is to reduce the MSE to obtain the optimum weights (W_*i*_s). These weights are independent, and trained separately for each data node. The impact or change (*I*) inflicted by each of the weather variables on the flu counts is estimated using the following formula.

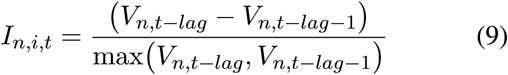 The impact value at node *n* coming from ith climatic variable at time *t* is the ratio of change happening before the appropriate situational time-lag (lag) from the time step *t* to the actual numeric data, *V*, (not the symbolic data) of *i*th weather variable. An appropriate lag value is retrieved from the time lags computed in the previous step based on the flu count symbol at this time-step *t*.
3. *Spatio-temporal adjustment factor*: Geographical proximity, in general, has a strong effect on the influenza outbreak in a particular region. One can observe similar flu trends between data nodes that are in spatial proximity (as shown in Fig. 5) for both GTF and CDC data. This impact is captured by computing an adjustment factor from the nearby data nodes. Similar to the weather variables, each neighboring data node impacts on this data node independently from the other neighboring data nodes. Thus, a weighted summation of individual adjustment factors is used. Here, Widrow-Hoff learning [55] is used to train those weights. Similar to the impact weights, the mean square error (MSE) training criterion is used as the cost function. An adjustment factor coming from each neighboring node is the average of flu variation difference during the previous three time stamps at that node. The adjustment factor, *γ*, to be applied at data node *n* on the initial forecast at time step *t* is the average of changes in the flu counts obtained at other nearby data nodes at time step *t* − 1.

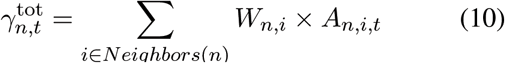 Total adjustment *γ*_*n*,*t*_ at data node *n* and time *t* is the average weighted summation of the individual adjustments *A*_*n*,*i*,*t*_ coming from all its neighbors that are in geographical proximity of *n*. Similar to the impact weights, adjustment weights (*W*_*n*,*i*_) are also trained using the Widrow-Hoff algorithm on the historical data from this node as well as its neighbors.

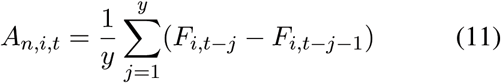 Individual adjustment (*A*_*n*,*i*,*t*_) for the neighbor *i* to data node *n* at time *t* is the average change in the previous *y* time steps. Here *F*_*i*,*t*−*j*_ is the actual flu count at neighbor *i* to *n* at time *t* − *j*. In our experiments we selected *y* to be 3 as it gave us optimal results.
4. *Forecast value estimation*: The total impact, defined in Eq. 8, is applied to the forecast value predicted by the LSTM (*F*^LSTM^), to calculate initial forecast, *F*^ini^.

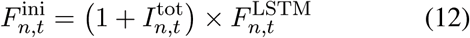 Final forecast after applying adjustment factor *γ*_*n*,*t*_ as computed in Eq. 10, *F*^final^, of data node *n* at time *t*

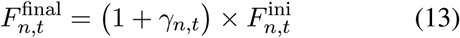

**Fig. 1:**
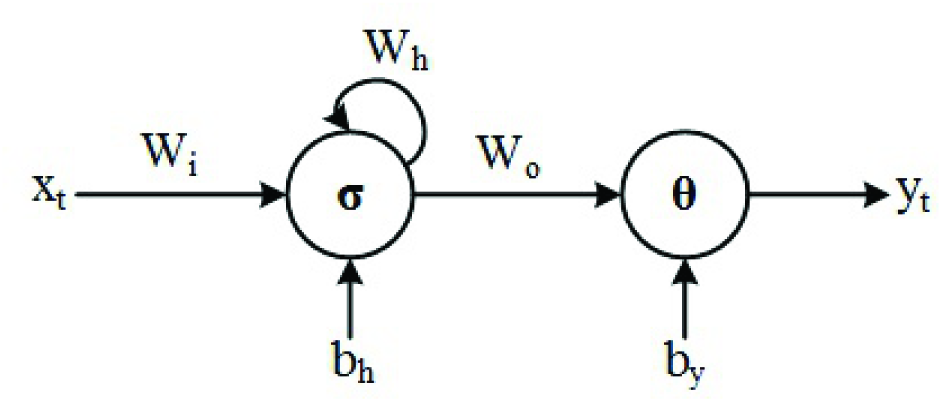
**Recurrent neural network**.

**Fig. 2:**
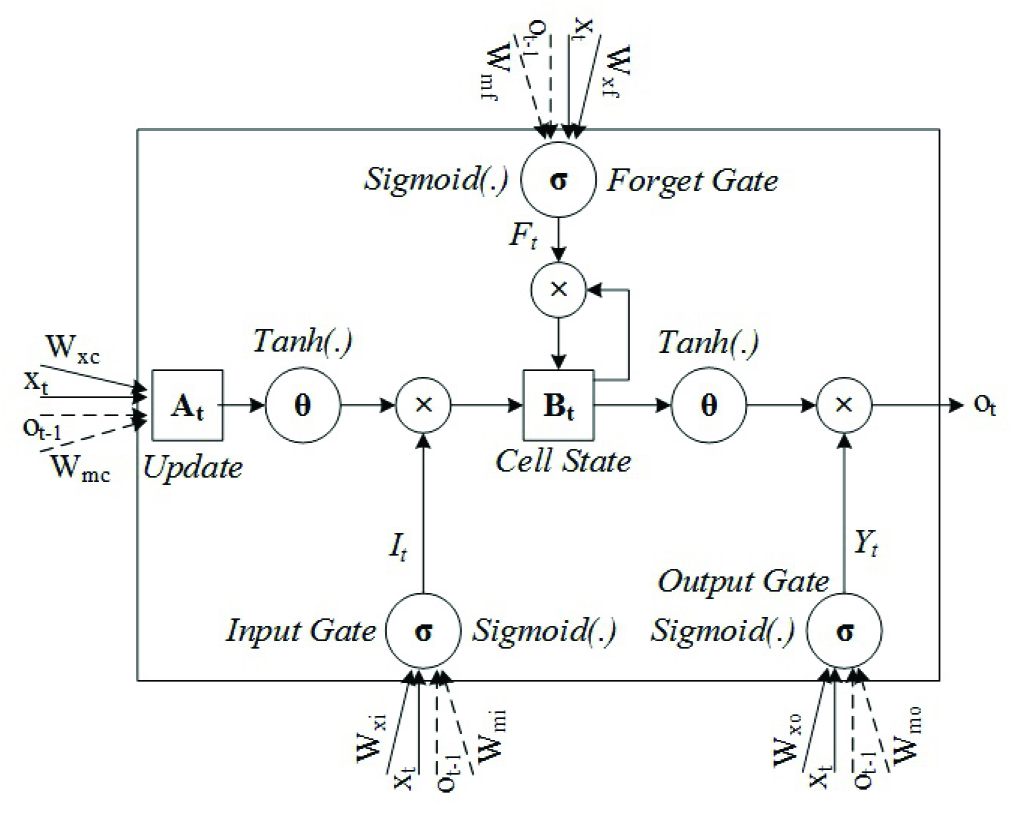
An LSTM cell containing the input gate, the forget gate, and the output gate. Each gate receives two vectors as input: x_*t*_, and previous output, o_*t−1*_.

**Fig. 3:**
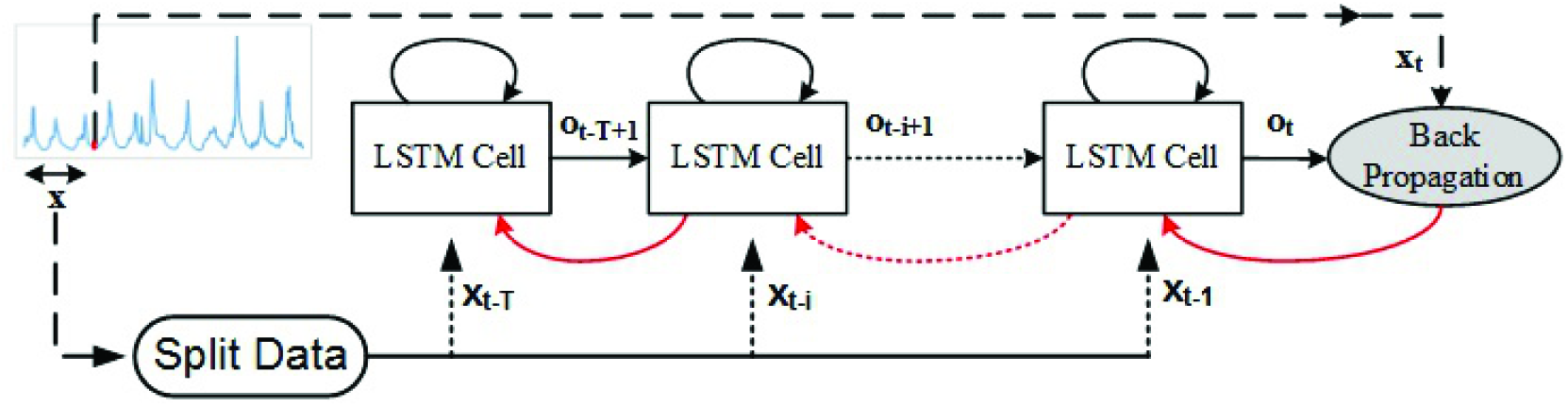
LSTM neural network consisting of the unrolled LSTM cells. The red backward arrows show the backpropagation algorithm and are not part of the network architecture.

**Fig. 4:**
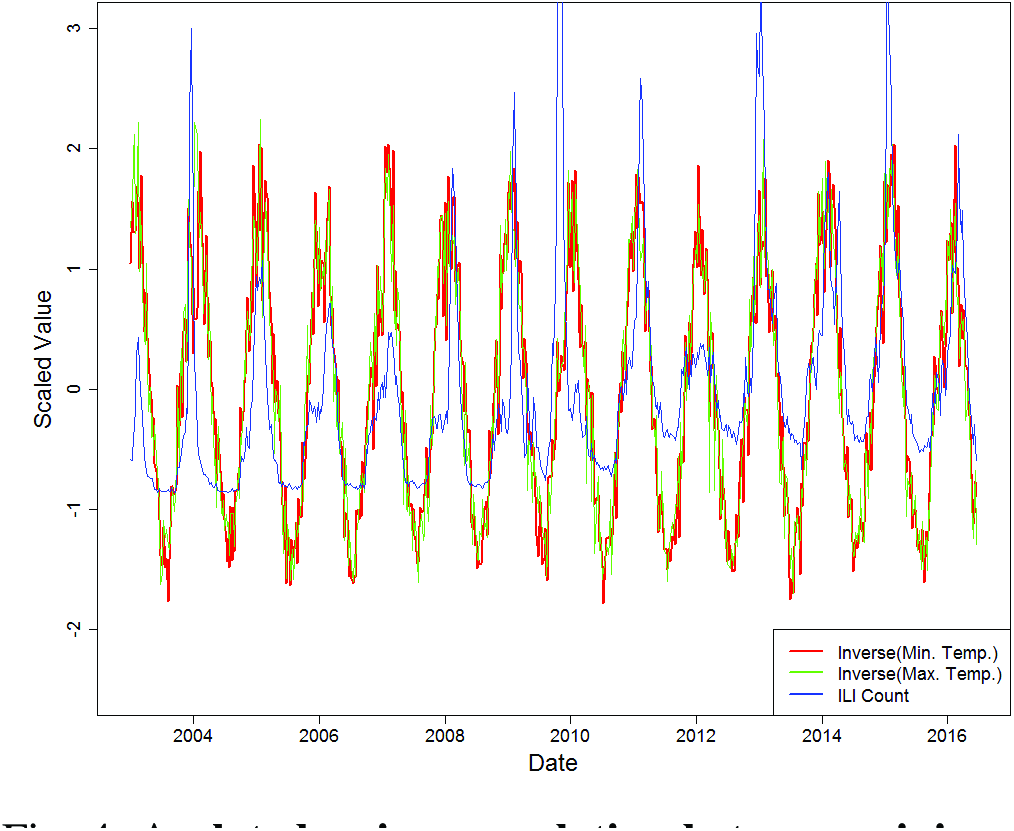
**A plot showing correlation between minimum and maximum temperatures and flu counts**.

**Fig. 5:**
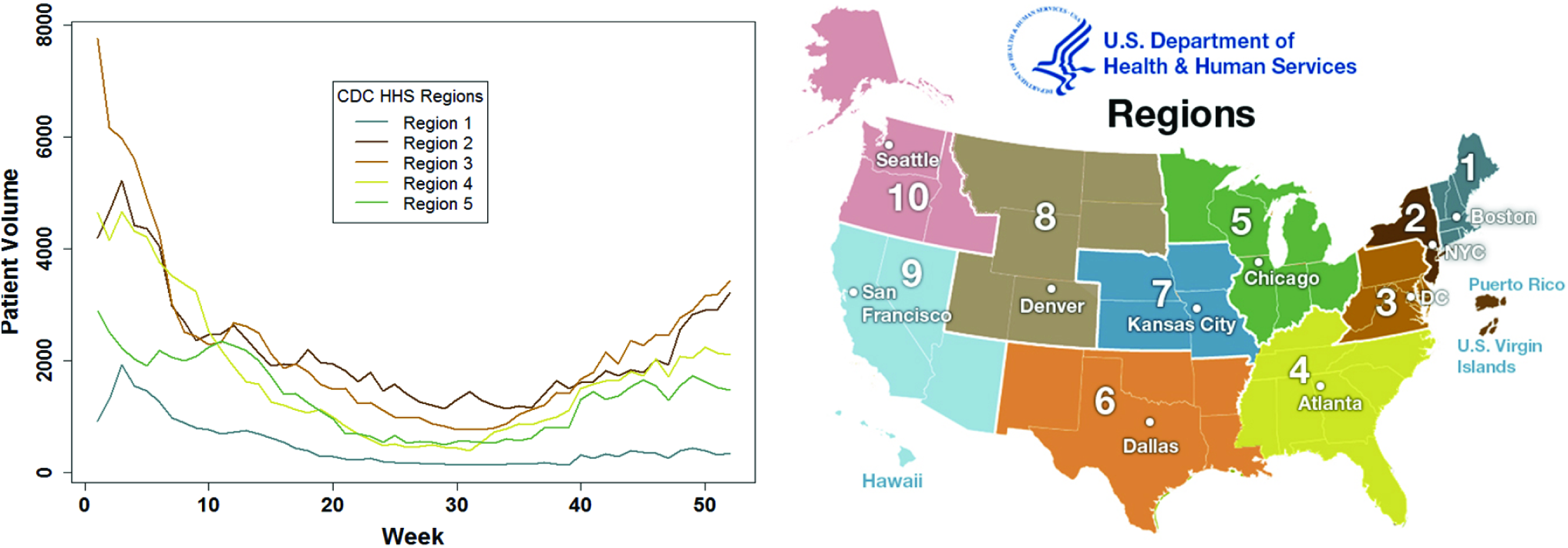
Flu count trends. A plot showing similar trends in flu counts in 2015 for different CDC regions (left). A map showing the CDC-HHS regions (right).

## III. Results

The three proposed data-driven models (LSTM, LSTM+CI, and LSTM+CI+SA) are compared with three ARIMA based models (ARIMA, ARIMA+CI and ARIMA+CI+SA) on two different publicly available data sets related to influenza counts, namely the CDC and GFT data sets. Both of these data sets represent a very broad sample in terms of spatio-temporal granularity. The models were evaluated by randomly generating 5 different samples from historical influenza counts. Each sample selected in the experiment represents a time-step (start week) from history that includes the training set and 20 time steps that are the testing set. The samples are selected in such a way that the 20 weeks to be forecast do not overlap with the other 4 experiments along this dataset. In other words, the validation sets are separately selected. The data between 1997-2014 and 2004-2013 were used for training the CDC and GFT data sets, respectively. The model was evaluated on two widely accepted evaluation metrics: the Mean Absolute Percentage Error (MAPE) and the Root Mean Square Error (RMSE). These were used in [56], [57]. All of the implementation of various models was done in R [58]. The LSTM model was implemented using the Tensorflow library [59].

### A. Evaluation criteria

The prediction performance of the proposed system is evaluated using the following metrics:

Mean absolute percentage error (MAPE) measures the average percent of absolute deviation between actual and forecasted values.

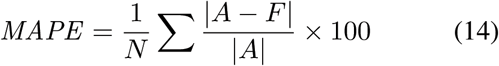

Root mean squared error (RMSE) captures the square root of average of squares of the difference between actual and forecasted values.

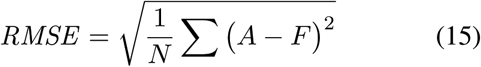

Root mean squared percentage error (RMSPE) captures percentage of square root of average of squares of the deviation between actual and forecasted values.

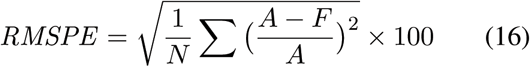

where, *N* is the number of test samples, *A* is the actual flu count, and *F* is its respective forecasted value.

We compared our results with the state-of-the-art ARIMA method. We also compared the results of our model at different phases; (i.e. LSTM prediction vs. LSTM with climatic variable impact vs. LSTM with climatic variable impact and spatial adjustment factor). We also tried to apply our climatic variable impact and spatial adjustment factor on top of ARIMA to evaluate their effectiveness. In experiments, we developed six models, composed of LSTM, ARIMA, climatic variable impact (CI), and spatial adjustment factor (SA), as follow:

- LSTM (The value predicted by LSTM (*F*^LSTM^) alone, that is without the variable impact or adjustment factor applied to it.)
- LSTM+CI(The estimated value after climatic variable impact factor is applied but not the spatial adjustment factor 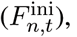 as computed in Eq. 12.)
- LSTM+CI+SA (This is the final forecast value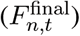 after both climatic variable impact factor and spatio-temporal adjustment factor are added to LSTM, as computed in Eq. 13. This is the proposed approach.)
- ARIMA (Flu count estimated using the state-of-art ARIMA.)
- ARIMA+CI (Flu count after climatic variable impact factor computed in Eq. 8 is applied to the simple ARIMA forecast.)
- ARIMA+CI+SA (Flu count after climatic variable impact factor as in Eq. 8 and spatio-temporal adjustment factor as in Eq. 10 are added to the simple ARIMA forecast.)

### B. Results for the CDC Dataset

Table I shows the comparison of the 6 forecasting models when these models were applied on all the ten geographical regions from HHS. The table compares the prediction performance of the selected models upto 20 weeks into the future. As mentioned earlier in this section, at each data node 5 random experiments were done making it 50 experiments overall for this dataset (5 experiments at each of the 10 CDC regions). The average performances of the proposed models in terms of the MAPE, RMSPE, and RMSE (% ILI) are shown in Table I. The LTSM model has the minimum MAPE, RMSPE, and RMSE (% ILI) when compared to ARIMA model. This itself is a significant improvement in forecasting accuracy. By integrating the climatic and spatio-temporal components into the LTSM model, we observe further improvement in forecasting performance. We also observe that by adding the climatic and environmental components to the ARIMA model, while the 1 week ahead forecast does not show any significant improvements, the 5 to 15 week forecasts have better performance compared to the baseline ARIMA model.

**Table I.** **ILI count predicted over 1, 5, 10 and 15 weeks using proposed models and ARIMA for the CDC dataset.**

| Weeks | 1-week |  |  | 5-weeks |  |  | 10-weeks |  |  | 15-weeks |  |  |
| --- | --- | --- | --- | --- | --- | --- | --- | --- | --- | --- | --- | --- |
| MODEL | MAPE | RMSPE | RMSE | MAPE | RMSPE | RMSE | MAPE | RMSPE | RMSE | MAPE | RMSPE | RMSE |
| LSTM | 21.38 | 29.31 | 0.26 | 57.09 | 80.66 | 0.58 | 62.32 | 78.82 | 1.59 | 70.05 | 96.18 | 1.46 |
| LSTM+CI | 21.13 | 29.17 | 0.25 | 57.1 | 80.96 | 0.58 | 62.2 | 78.58 | 1.61 | 69.67 | 94.76 | 1.46 |
| *LSTM+CI+SA* | *16.69* | *23.13* | *0.22* | *51.49* | *72.58* | *0.55* | *60.47* | *76.28* | *1.54* | *65.86* | *87.93* | *1.41* |
| ARIMA | 44.69 | 83.58 | 0.3 | 68.99 | 95.75 | 0.68 | 79.89 | 100.46 | 1.78 | 109.6 | 154.32 | 1.93 |
| ARIMA+CI | 45.2 | 83.5 | 0.3 | 69.11 | 95.51 | 0.68 | 80.06 | 100.74 | 1.76 | 110.85 | 156.85 | 1.94 |
| ARIMA+CI+SA | 45.73 | 86.02 | 0.28 | 62.03 | 85.25 | 0.67 | 77.82 | 97.76 | 1.71 | 103.15 | 143.13 | 1.88 |

Fig 6 shows 9 charts that compare all 6 models for 3 HHS regions, namely Region-2 (Row A), Region-5 (Row B) and Region-10 (Row C) with respect to MAPE, RMSPE and RMSE (% ILI). Fig. 7 shows the actual and predicted ILI counts for those three regions from one of the test samples. It can be seen that the numbers from Table I correlate with the plots from Fig 6, LSTM and its variants outperforming ARIMA and its variants in most of the cases. For Region-10 (plot from Row C of Fig 6), in the later weeks (weeks 15 to 20) of forecasting ARIMA performs better than LSTM. Fig. 7 also shows that LSTM and its variants are able to follow the actual ILI counts trend line during the first 5 weeks; this might be because of the proposed model being able to capture the impacts from climate variables and spatio-temporal factors accurately. Additionally, both the table and plots show the importance of impact components used by the proposed models to significantly reduce the error.

**Fig. 6:**
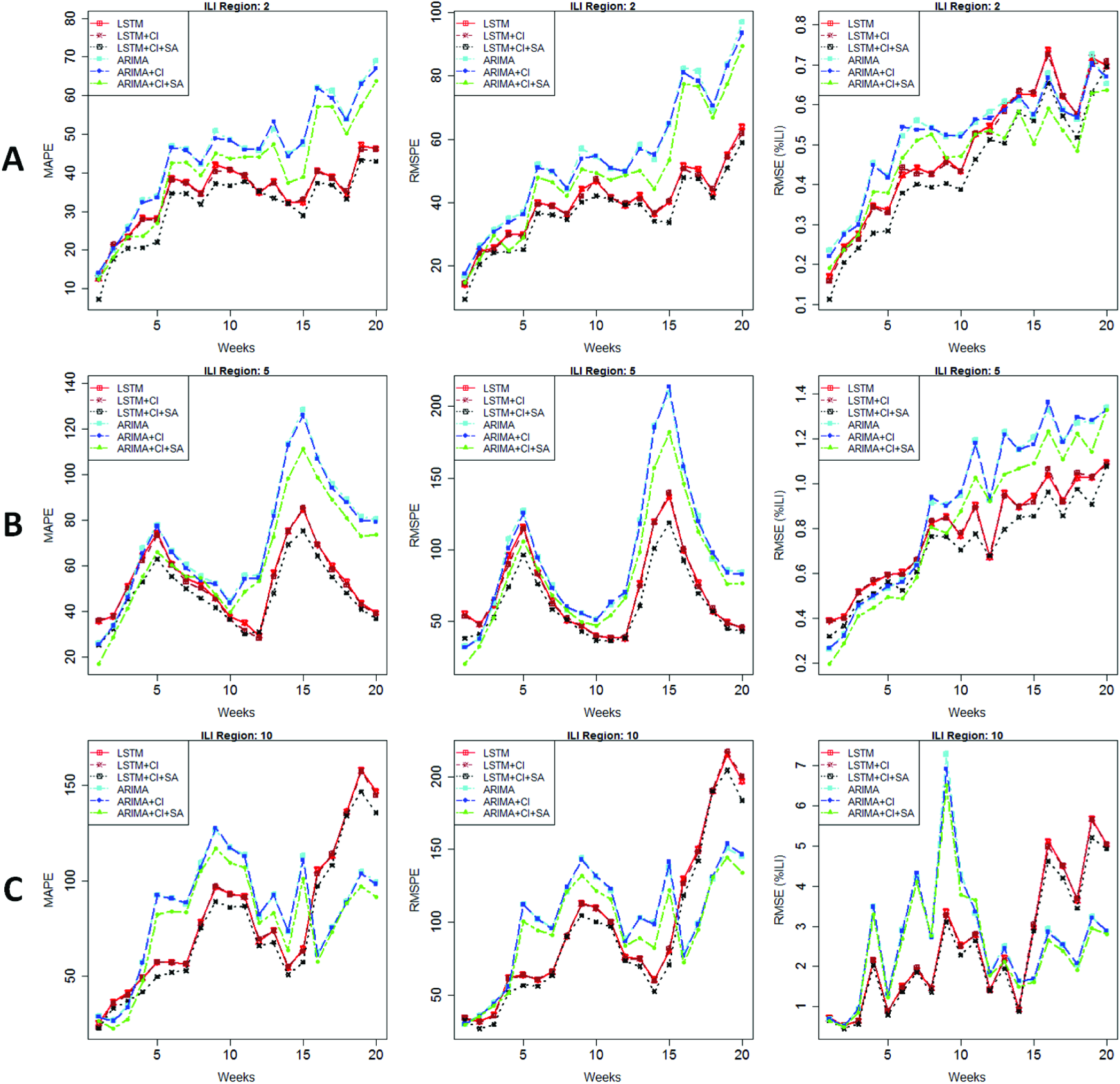
MAPE, RMSPE and RMSE (%ILI) of the flu prediction models over 20 weeks applied on ILI count of CDC dataset. A: Region 2, B: Region 5, and C: Region 10.

**Fig. 7:**
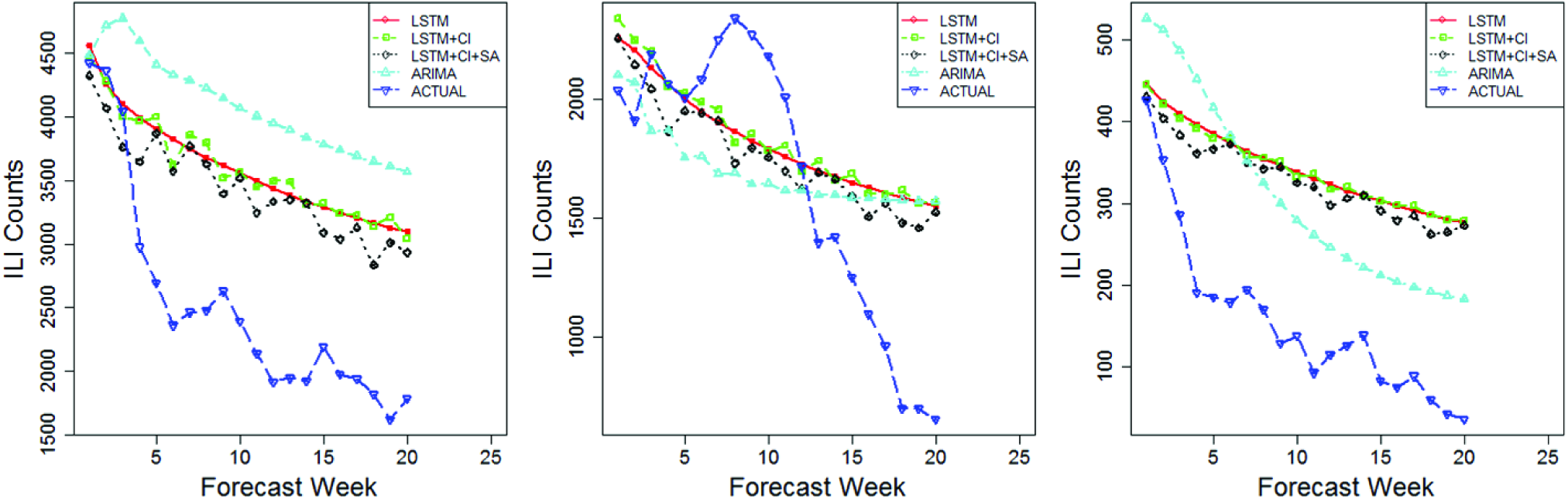
**Actual and predicted ILI count for regions 2, 5, and 10 from left to right respectively**.

### C. Results for the GFT dataset

Similar to the CDC results, Table II shows the comparison of the forecasting models discussed earlier for the 6 cities from Google Flu Trends data. Again, 5 different experiments were conducted at each city, thereby creating 30 test samples at each time step. The same error metrics MAPE, RMSPE, and RMSE (% ILI) are used to evaluate the 6 models. The proposed approach (LSTM + CI + SA) is better than the other 5 models compared in our analysis. Overall the LSTM and its variants are more accurate than the ARIMA and its variants. It can also be seen that impact component from climate variables is adding noticeable improvement to the base LSTM and ARIMA models, but the spatio-temporal component is improving the accuracy of these base models significantly. Unlike in CDC results, the magnitude of these errors is much less with the GFT datasets, because of the smoothness and high volumes in GFT data.

**Table II.**
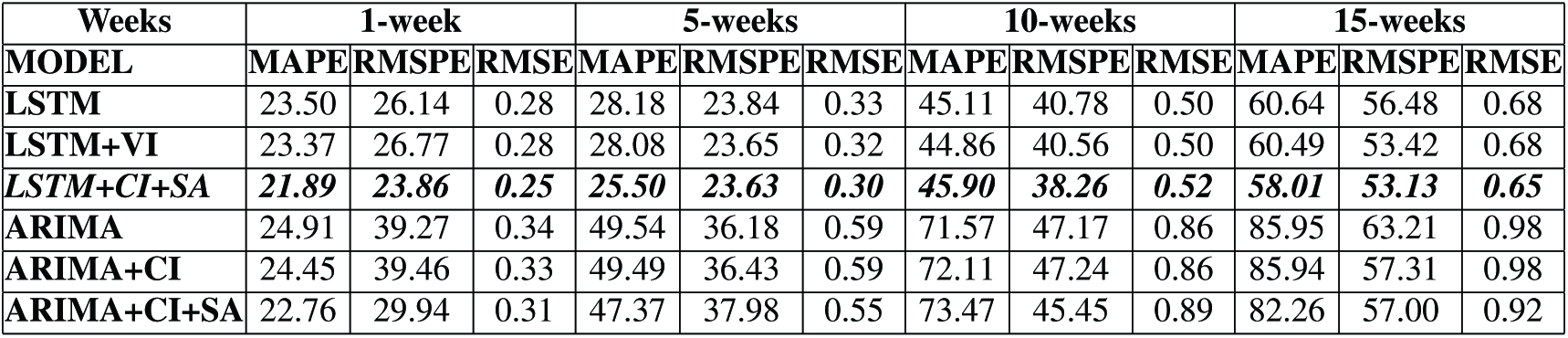
**Flu count predicted over 1, 5, 10 and 15 weeks using proposed models and ARIMA for the GFT dataset**.

Fig. 8 shows the error charts of MAPE, RMSPE, and RMSE (% ILI) for three cities Austin (row A), Dallas (row B), New Orleans (row C) and Fig. 9 shows the comparison of the predicted values from these models with the actual GFT volumes. Compared to CDC dataset, the proposed approach is much better and outperforms ARIMA significantly all along 20 weeks of prediction and across all cities. Similarly, the addition of the impact components from spatio-temporal and climate variables improves the performance of both ARIMA and LSTM base models. Fig. 9 also demonstrates that the proposed models show strong correlation with actual data, at least until 12 weeks.

**Fig. 8:**
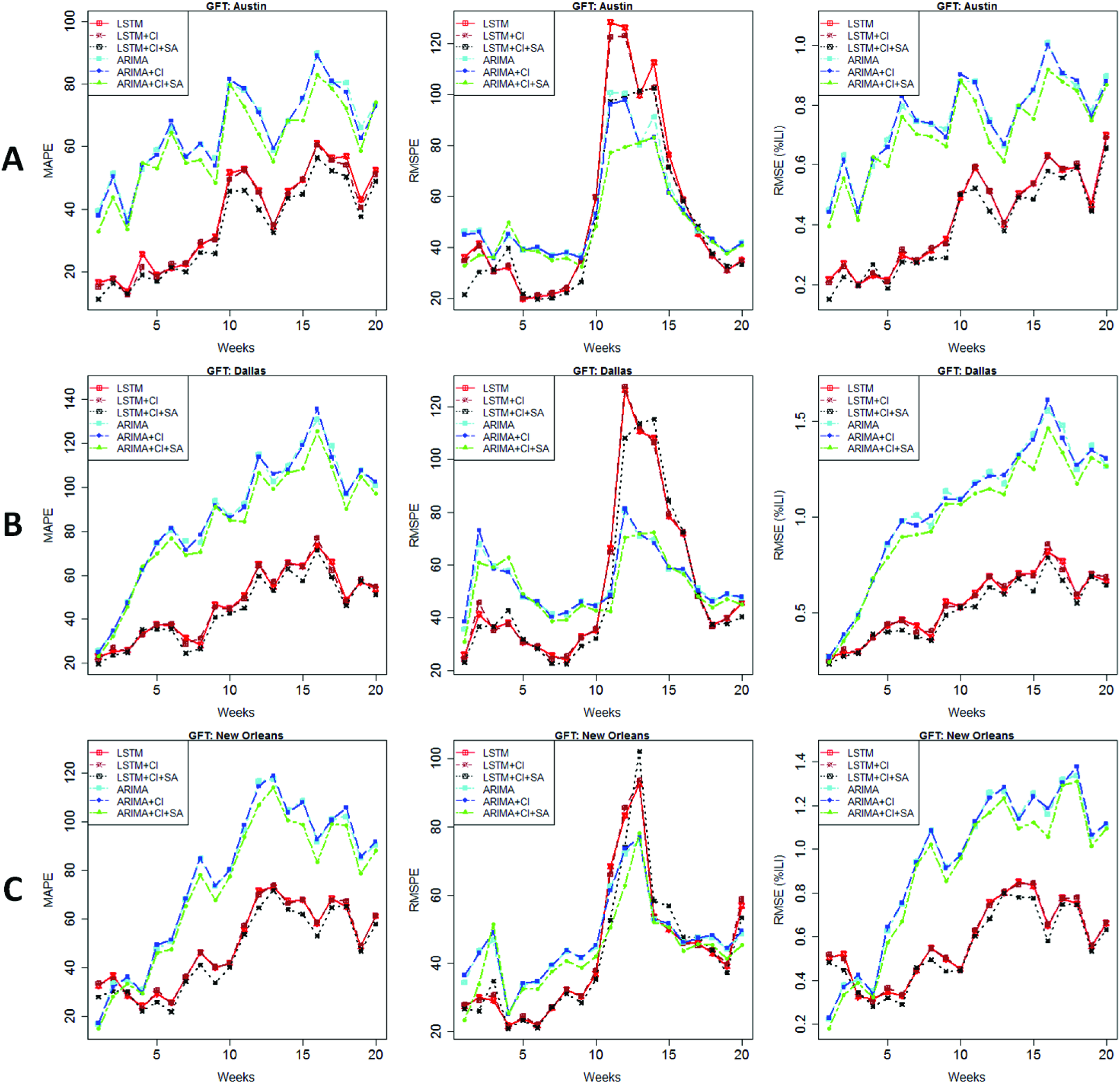
Fig. 8: MAPE, RMSEP and RMSE (%ILI) of the flu prediction models over 20 weeks applied on the GFT dataset. Austin, Dallas, and New Orleans are randomly selected.

**Fig. 9:**
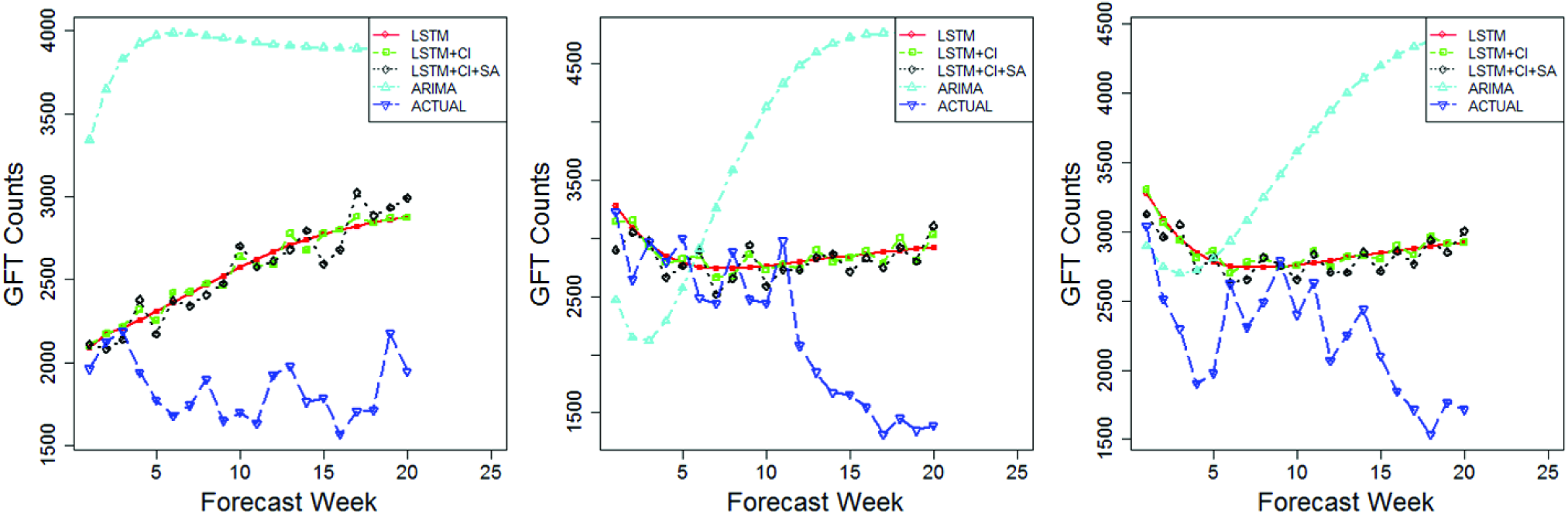
**Actual and predicted GFT for Austin, Dallas, and New Orleans from left to right respectively.**

## IV. Conclusion

In this paper, we proposed data driven approaches to improve influenza forecasting. The first key contribution is the applicability of the LTSM based deep-learning method which is shown to perform well compared to existing time series forecasting methods. We further reduced the error of the deep learning based forecasting method by introducing an approach to integrate the impacts from climatic variables and spatio-temporal factors. We evaluated the proposed approach on publicly available CDC-HHS ILI and GFT datasets. The results also showed that the impact component integrated into the baseline models (LSTM and ARIMA) significantly improved their performances. The proposed method offers a promising direction to improve the performance of real-time influenza forecasting models. Additionally, the proposed method may be useful for other serious viral illnesses such as Ebola and Zika.

In this paper, we have implemented separate learning components for the climatic variables and for the geospatially proximal variables. Our future study seeks to develop an end-to-end learning model incorporating all the modules. This could be done by using a convolutional LSTM [60] to learn spatio-temporal patterns.

## Acknowledgment

This project is granted by:

- NAME: Division of Computer and Network Systems (CNS). Grant No: 1650551. URL: https://www.nsf.gov/div/index.jsp?div=CNS.
- NAME: Oak Ridge Associated Universities (ORAS). Grant No: 370270. URL: https://www.orau.org/

The funders had no role in study design, data collection and analysis, decision to publish, or preparation of the manuscript.

